# Using split protein reassembly strategy to control PLD enzymatic activity

**DOI:** 10.1101/2024.01.27.577557

**Authors:** Yuanfa Yao, Changyi Hu, Li Jianxu, Xiayan Lou, Gaojun Chen, Xiaohan Qian, Lian He, Xuesong Li, Peng Zhang, Yingke Xu, Hanbing Li

## Abstract

Phospholipase D (PLD) and phosphatidic acid (PA) play a spatio-temporal role in regulating diverse cellular activities. Although current methodologies enable optical control of the subcellular localization of PLD and by which influence local PLD enzyme activity, the overexpression of PLD elevates the basal PLD enzyme activity and further leads to increased PA levels in cells. In this study, we employed a split protein reassembly strategy and optogenetic techniques to modify superPLD (developed by Jeremy Baskin group). We splited this variants into two HKD domains and fused these domains with optogenetic and chemogenetic elements and by which we achieved control of the two HKD interaction and then restored the PLD enzymatic activity.

---

Phospholipase D (PLD) and phosphatidic acid (PA) have been shown to play crucial roles in diverse cellular physiological and pathological processes ^1^. However, the spatiotemporal regulatory mechanisms of both remain largely unknown in cells. To address this, there is a critical need for a highly efficient platform capable of activating PLD and detecting PA. The enzymatic activity of cellular PLD and the levels of intracellular PA are notably low under non-stimulated conditions. PLDs have been demonstrated to be activated by indirect agonists, including PMA, EGF, and small G proteins ARF1/6, RHOA, ARL11/14 ^2^. To directly increase the PLD activity in subcellular structure, Baskin et al. employed optogenetic techniques to manipulate the subcellular localization of bacterial PLD_PMF_ for PA production ^3^. Their recent publication reported an ingenious PLD construction to control its activity by inserting optogenetic element LOV2 into a highly-active PLD_PMF_ variant ^4, 5^. Here, we propose a different strategy for control PLD activity with optogenetic or chemogenetic elements through a split protein reassembly.

Before this, we also attempted to control the activity of PLD_α_1, a PLD isoform with relatively high basal enzyme activity, through optogenetic element LOV2 domain. We inserted the LOV2 domain near the unique lid structure of the substrate pocket of PLD_α_1 ^6, 7^. We hypothesized that the light-induced unwinding of LOV2 would alter the conformation of the lid, thereby regulating the activity of PLD_α_1. However, our results showed blue light stimulation abolished the total PLD activity in cells with stable expression of LOV2-inserted PLD_α_1 (LOV2 at residue Thr277 site) (**Fig. S1A-C**). Given that the C2 domain is required for PLD_α_1 enzyme activity, we subsequently hypothesized that through splitting PLD_α_1 at the linker between the C2 domain and the catalytic domain (PLDCTD), we might be able to control the dimerization of C2 and PLDCTD domain based on optogenetic and consequently restoring the enzyme activity of the split PLD_α_1. We utilized the iLID system to fuse corresponding split PLD_α_1 fragments, namely Tom20-C2-LOV2 (SsrA) and mCh-SspB-PLDCTD. The AlphaFold2 tool was employed to predict the assembled structure and assess the spatial accessibility of C2 and PLDCTD for reassembly. Upon co-expression of both constructs in HeLa cells, dynamic changes in the localization of PLDCTD to mitochondria were observed upon blue light stimulation (**Fig. S1D-G**). Despite the successful dimerization of the C2 and PLDCTD domains, enzymatic activity assays indicated no significant increase in total cellular PLD activity (**Fig. S1H**). This outcome suggests that the splited PLD_α_1 activity was not restored. Furthermore, we noted the spontaneous dimerization of the two HKD domains of human PLD1 when we splited human PLD1 into two parts at Ser 584 residue (**Fig. S1I**). This finding suggests that both plant PLD_α_1 and human PLD1 may not be a suitable candidate for optogenetic modification.

Very recently, Baskin et al. undertook efforts to optimize PLD_PMF_ activities through directed evolution, leveraging their ingenious phosphatidic acid (PA) detection technique IMPACT ^5^. They successfully generated a range of PLD_PMF_ variants exhibiting significantly enhanced enzymatic activity. Notably, certain variants demonstrated a remarkable 100-fold increase in activity, referred to as superPLD. Hence, we selected superPLD as the target for our optogenetic and chemogenetic modification. To identify potential splitting positions, we employed a flexible, long linker to connect two splited subunits. We observed that superPLD with insertions of the linker at Pro 285 and Ala 290 residues maintained enzyme activity, resulting in the accumulation of PA probes Opto-PASS (a improved PA biosensor with optogenetic^8^) on mitochondria (**Fig. 1A-B**).

**Figure 1.**
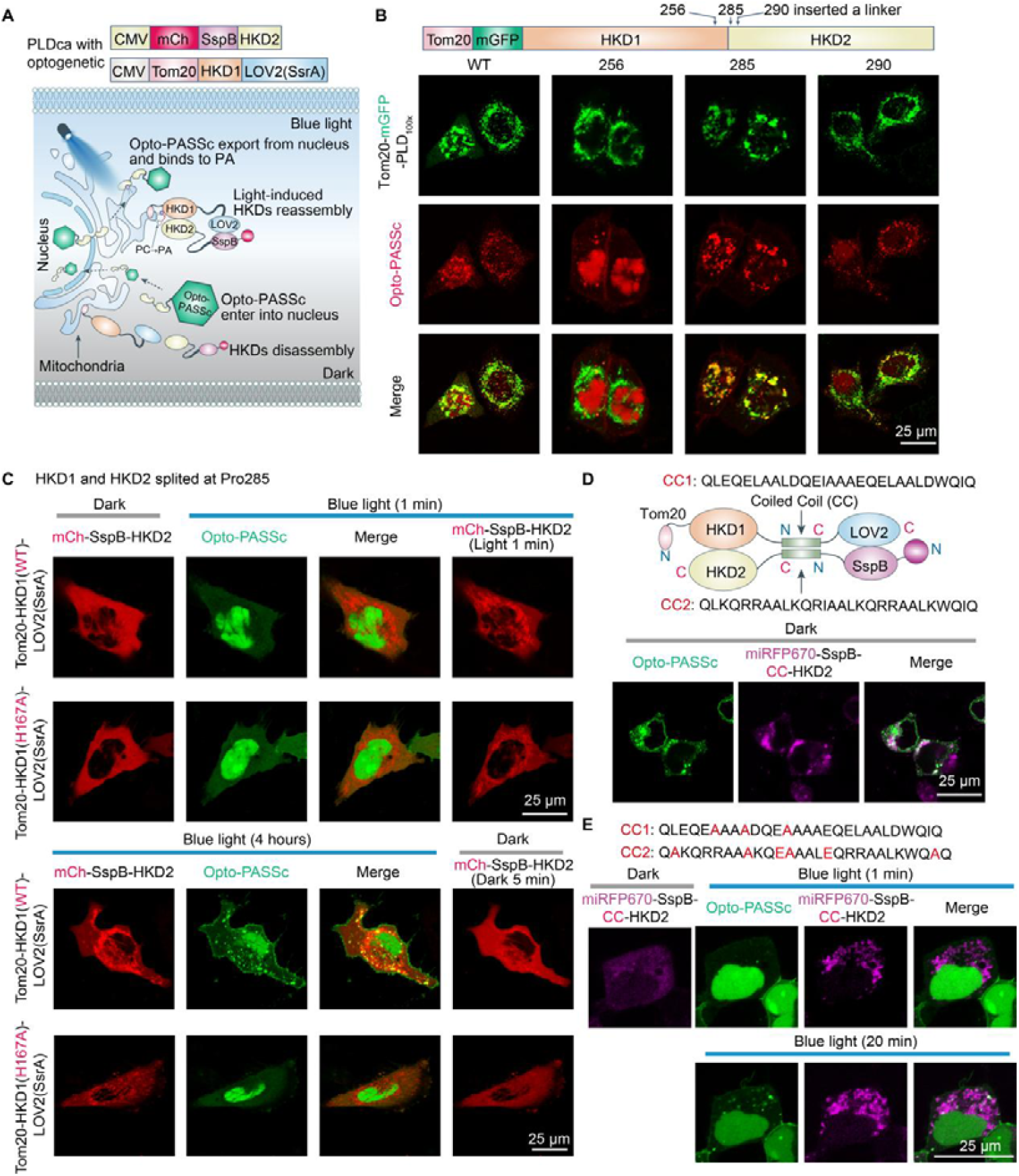
Using the split protein reassembly strategy to modify PLD and control its activity via light. A, the conceptual diagram of optogenetic control of superPLD enzyme activity. B, using PA probe Opto-PASSc to screen the split site of the superPLD by insertion of a linker. C, blue light stimulation induced the dimerization of splited PLD on mitochondria and restored its activity. D, the CC peptides were integrated into the iLID system to enhance interactions between HKD1 and HKD2. E,the incorporation of the optimized CC peptides allowed for the regulation of superPLD activity through light.

Subsequently, we splited superPLD at Pro 285 and Ala 290 residues and fused it with corresponding optogenetic tools, namely Tom20-HKD1(285/290)-LOV2(SsrA) and corresponding mCh (or miRFP670, here refered to miRFP670nano)-SspB-HKD2. Following blue light stimulation, we observed an increase in PA levels on mitochondria using the Opto-PASS probe, compared to the dead PLD variant containing the H167A mutation in HKD1 (**Fig. 1C** and **Fig. S2A**), which indicates dimerization of the two HKD domains of superPLD resulted in the restoration of its enzyme activity. However, the catalytic efficiency of this PLD variant remained limited, as increased mitochondrial PA was detected in only a subset of cells. To enhance efficiency, we shortened the flexible linkers between the HKD domains and the optogenetic elements. Unfortunately, these modifications did not yield the anticipated improvement (data not shown).

As the structural details of iLID after dimerization remain unresolved, we hypothesized that the low efficiency of our system might result from improper orientation of the HKD1 and HKD2 domains during dimerization. To address this, we introduced a pair of coiled-coil (CC) sequences consisting of antiparallel _α_-helices into the iLID system, termed here as iLIDc. This antiparallel CC subunit demonstrated a self-dimerization capability and by which aided HKD1 and HKD2 in facilitating optimal interactions (**Fig. 1D**). The results demonstrated that iLIDc significantly enhanced interactions between the HKD domains, as Opto-PASSc was relocated to mitochondria even in the absence of light stimulation (**Fig. 1D**), which indicates that this iLIDc was no longer light-dependent due to the robust spontaneous dimerization of the CC subunit. Therefore, we attempted to mutate the CC part to weaken its dimerization, thereby restoring the light-controlled functionality of the iLIDc. It has been demonstrated that CC peptides contain a heptad repeat pattern, hxxhcxc, comprising hydrophobic (h) and charged (c) amino acid residues. After we substituted leucine and isoleucine with the less hydrophobic alanine, we observed that fluorescently-labeled HKD2 domain exhibited less localization on the mitochondria compared to the wild-type CC peptide fusion system in the dark (**Fig. 1D-E**), indicating CC peptides-mediated dimerization was diminished. Furthermore, after continuous pulsed blue light stimulation for 20 minutes, we observed the relocation of numerous PA biosensors to the mitochondria, suggesting that the modified iLIDc can activate splited PLD enzyme activity through blue light stimulation (**Fig. 1E**). The results were consistent with our hypothesis that using iLID system to bring two HKD domains into close proximity and the antiparallel CC peptides would assist these two components to assemble into an active PLD in a suitable orientation. The modified iLIDc system may also have the potential to be applied in manipulating other proteins function through a similar split reassembly strategy.

It is well established that blue light has limited tissue penetration, which directly impacts the activation of optogenetic tools when expressed in deep tissue. Therefore, to create a tool system capable of regulating PLD enzyme activity in deep tissues, we aimed to modify superPLD using the classical chemogenetic system rapamycin-FKBP-FRB. Based on structural information indicating the proximity of the N-terminus of the FRB domain to the C-terminus of FKBP, we constructed Tom20-BFP-HKD1-FRB and the corresponding miRFP670-FKBP-HKD2 (**Fig. 2A** and **Fig. S2B**). Following a 40-minute incubation with 100 nM rapamycin, both the HKD2 domain and PA biosensors notably translocated to the mitochondria (**Fig. 2B**), suggesting that the chemical-induced dimerization of HKD1 and HKD2 successfully restored their phospholipase activity. In our previous study, we observed that the endogenous PLD exhibited basal activity, and a significant portion of the overexpressed PA biosensor PASS remained unbound^8^. To minimize the influence of unbound PA biosensors on PA detection performance, we developed an enhanced PA biosensor, Opto-PASSc. This biosensor sequesters unbound biosensors in the cell nucleus and releases them in response to an increase in PA levels in the cytosol. Therefore, we expressed HKD1 and HKD2 in the cytoplasm. Using the Opto-PASSc biosensor, we visually observed a decrease in the nuclear signal but an increase in the cytosol following rapamycin treatment (**Fig. 2C**). This shift indicates the reactivation of superPLD activity and underscores the enhanced utility of Opto-PASSc as a PA-detecting tool. Furthermore, the results of PLD activities assessed using the IMPACT technique also illustrated that rapamycin-induced dimerization revived superPLD activity. Following the exclusion of the impact from endogenous PLD, a 7.5-fold increase was observed (**Fig.2D**).

**Figure 2.**
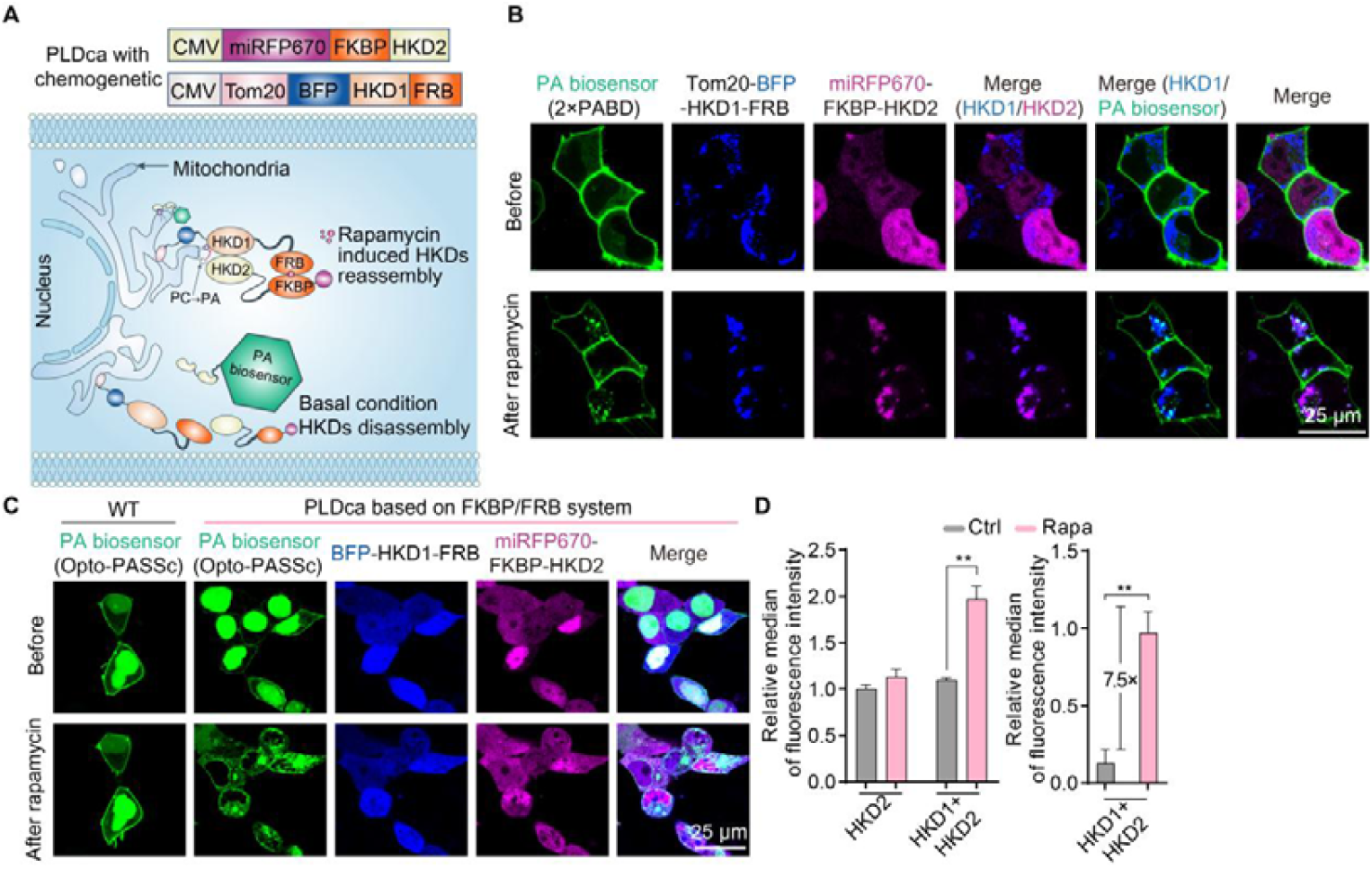
Using the split protein reassembly strategy to modify PLD and control its activity via chemogenetic system rapamycin-FKBP-FRB. A, the conceptual diagram of chemogenetic control of PLD enzyme activity. Using PA biosensor PASS (B) and Opto-PASSc (C) to assess the PLD activity change after rapamycin treatment. D, the IMPACT technique was employed to assess the change in PLD activity following rapamycin treatment, with single expression of miRPF670-FKBP-HKD2 serving as a negative control.

Interestingly, in cells with overexpression of superPLD, we noted a significant modification in mitochondrial morphology, characterized by a punctate structure. This alteration persisted even in the absence of anchoring superPLD to mitochondria through a mitochondrial targeting sequence (**Fig. S3A-B**). In addition, some of these punctate mitochondria also co-localized with lysosomes (**Fig. S3C**). To exclude the possibility of mitochondrial autophagy in this subset, we employed the mitochondrial autophagy probe Mito-Rosella. We observed that the fluorescence signal of PHluorin on these punctate mitochondria persisted. Upon addition of the mitochondrial autophagy inducer CCCP, the fluorescence signal diminished (**Fig. S3D**). This suggests that the punctate mitochondria did not fuse with lysosomes, indicating a lack of mitophagy. The morphological changes, therefore, are likely attributed to elevated intracellular PA levels resulting from the overexpression of superPLD, but the regulatory mechanism remain unclear.

---

**Supplementary Figure 1.**
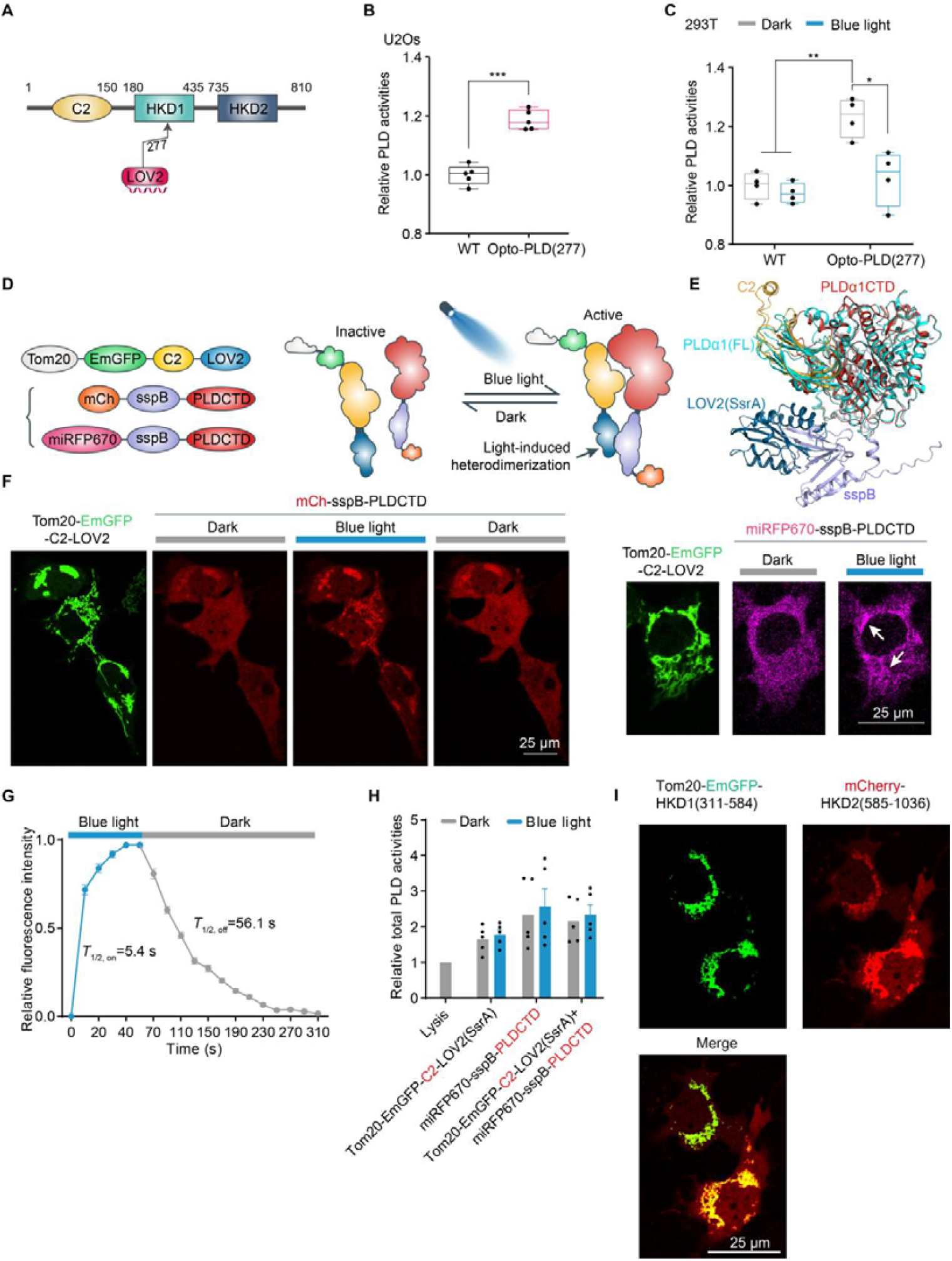
Optogenetic modification of Arabidopsis PLD_α_1. A-C, Insertion of LOV2 at Thr 277 residue caused an inhibitory effect of PLD_α_1 activity by blue light. D, the optogenetic design of split PLD_α_1 and its fusion with iLID system. E, the predicted secondary structure of Opto-PLD by AlphaFold2. F-G, the light-induced dimerization of PLDCTD and C2 domain and quantitative analysis of its dynamic. H, biochemical analysis of Opto-PLD enzyme activity in response to light stimulation. I, spontaneous dimerization of split human PLD1.

**Supplementary Figure 2.**
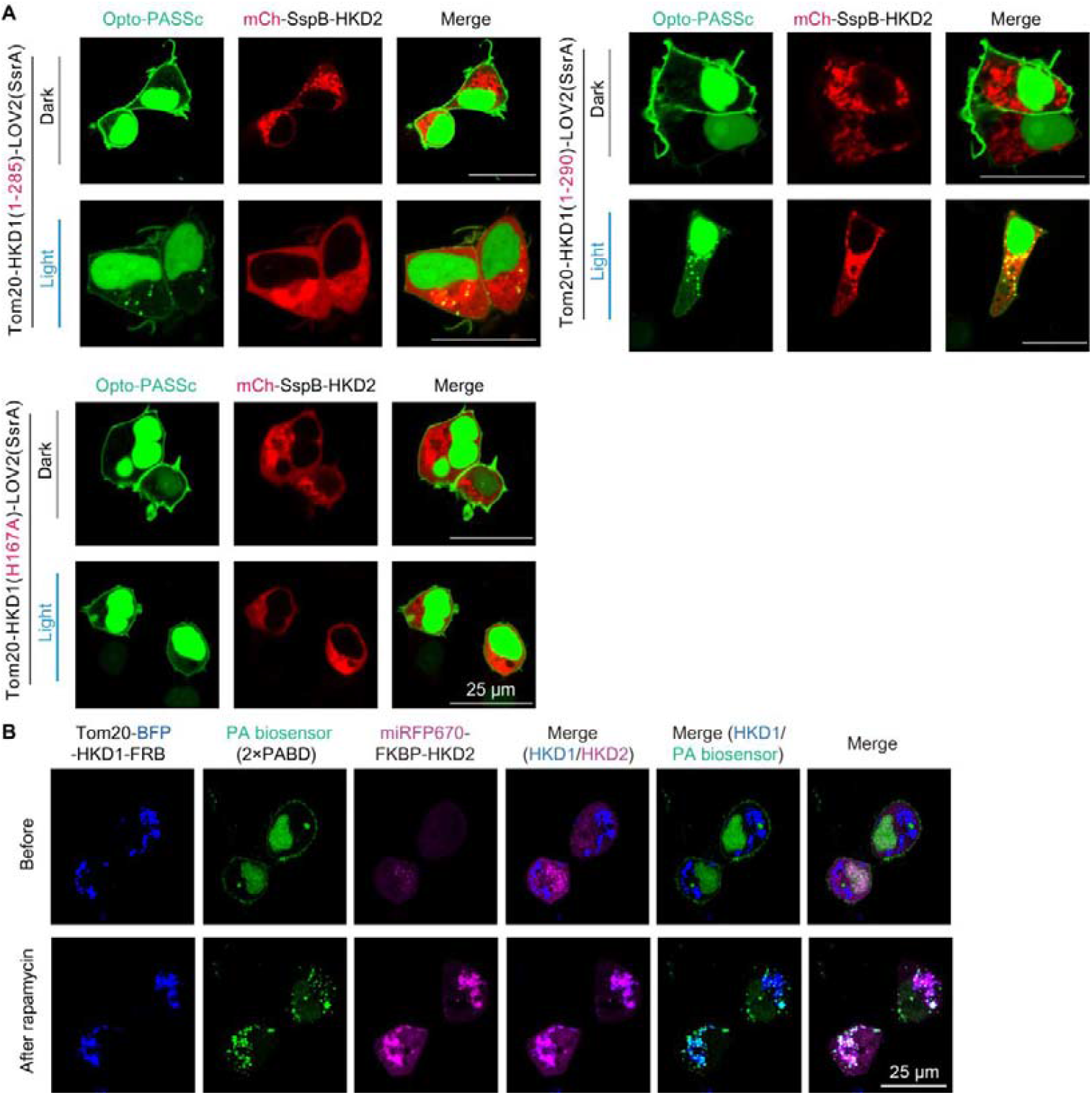
modifying superPLD based on optogenetic and chemogenetic system. A, Using the PA biosensor Opto-PASSc to evaluate the altered activity of split superPLD post light exposure, the split sites were identified as Pro 285 and Ala 290 residues. Additionally, the H167A variant was inactive in PLD activity. B, Using Opto-PASSc to assess the mitochondria-localized PLD activity before and after rapamycin incubation.

**Supplementary Figure 3.**
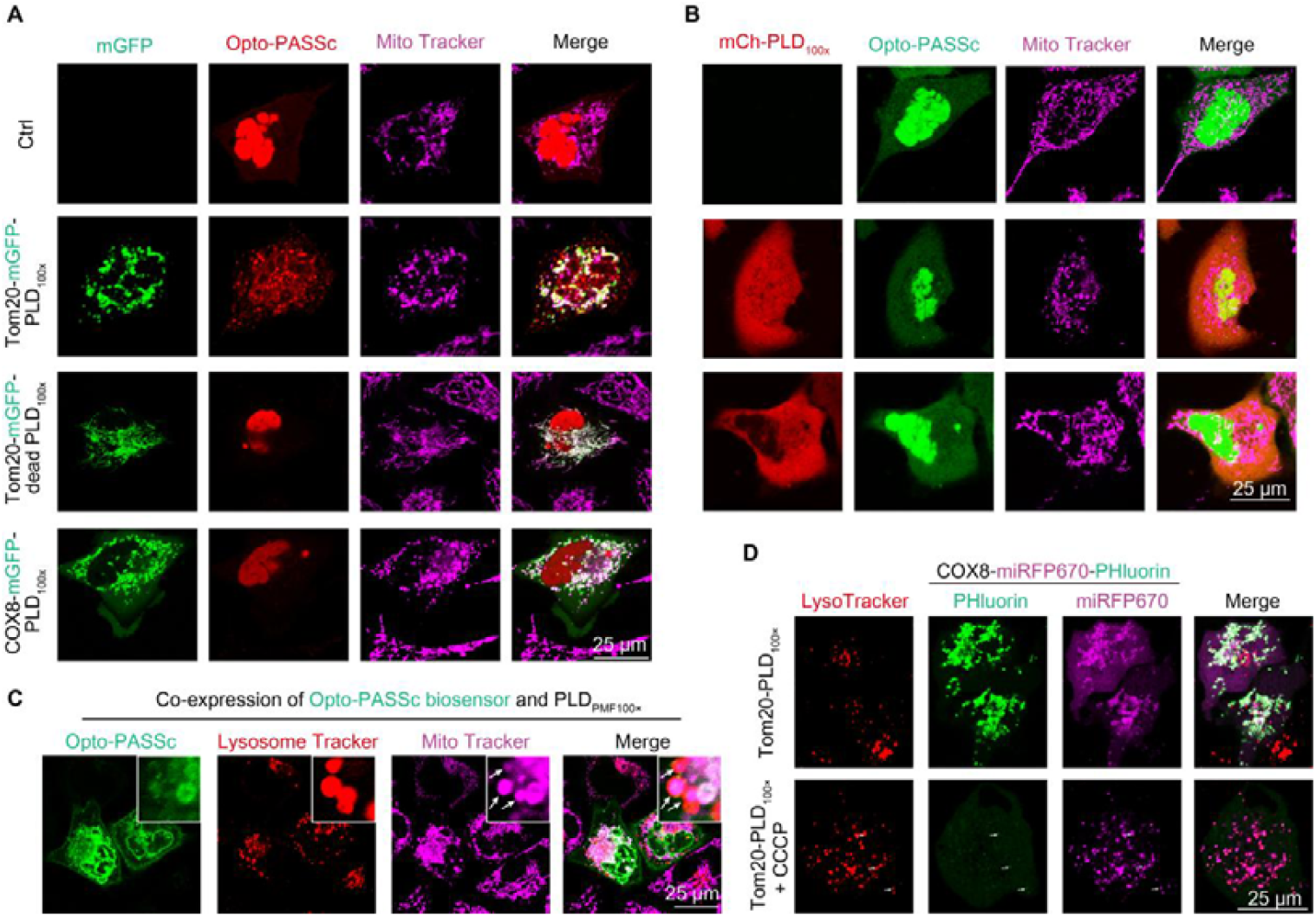
overexpression of superPLD altered the mitochondrial morphology. The mitochondria exhibited a punctate structure following the overexpression of superPLD in both the mitochondria (A) and the cytosol (B), PLD_100×_ here referred to superPLD. C, the co-localization of puncate mitochondria and lysosome after PLD overexpression. D, using Mito-Rosella (COX8-miRFP-PHluorin) to assess the mitophagy of the punctate mitochodria.

## Notes

### Competing Interest Statement

The authors have declared no competing interest.

### Summary of Updates

we have added new results in the revision and also revised the manuscript.

