## Supplementary figures and images for "Using split protein reassembly strategy to control PLD enzymatic activity"

### Supplementary figures.pdf

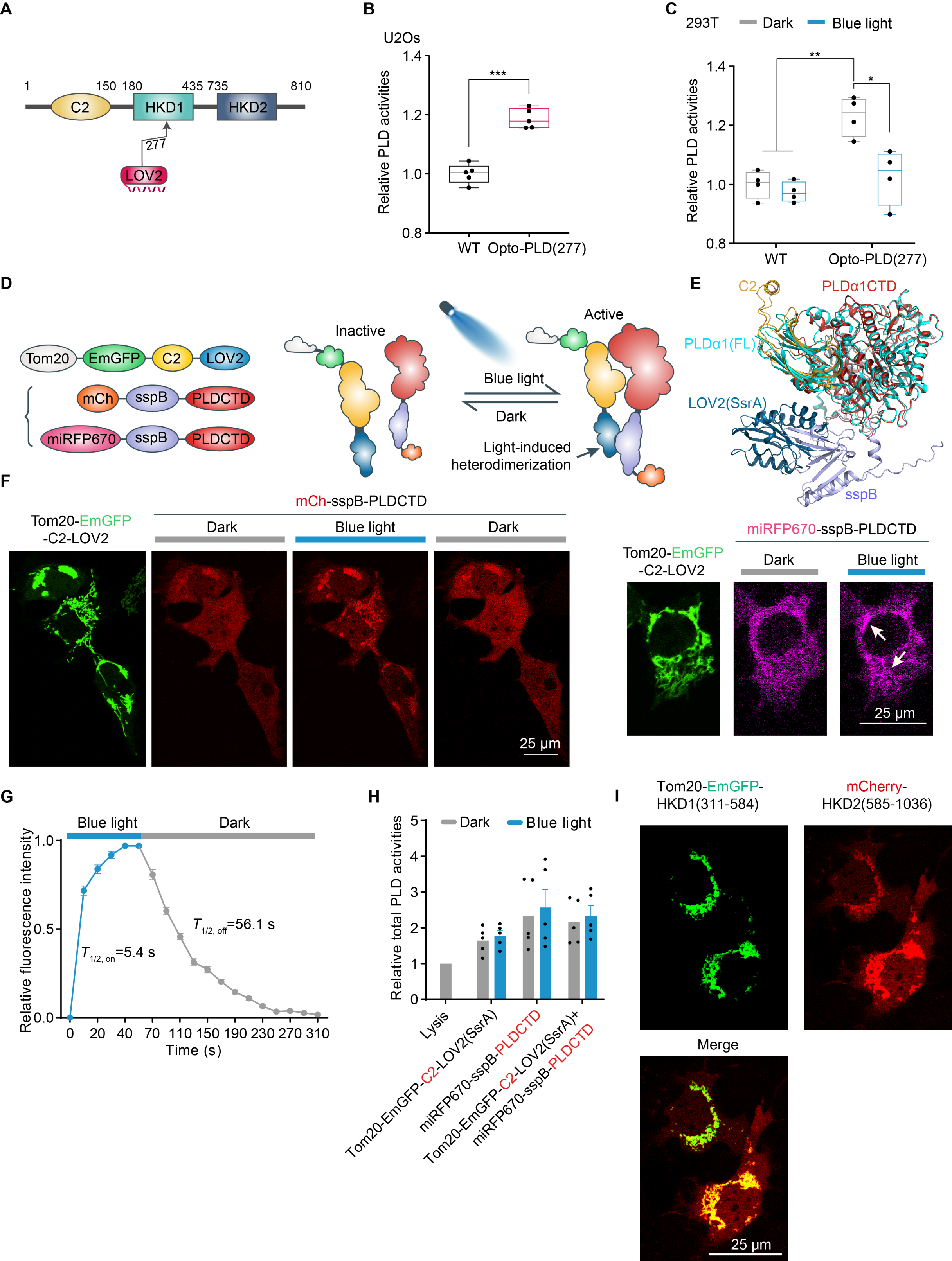

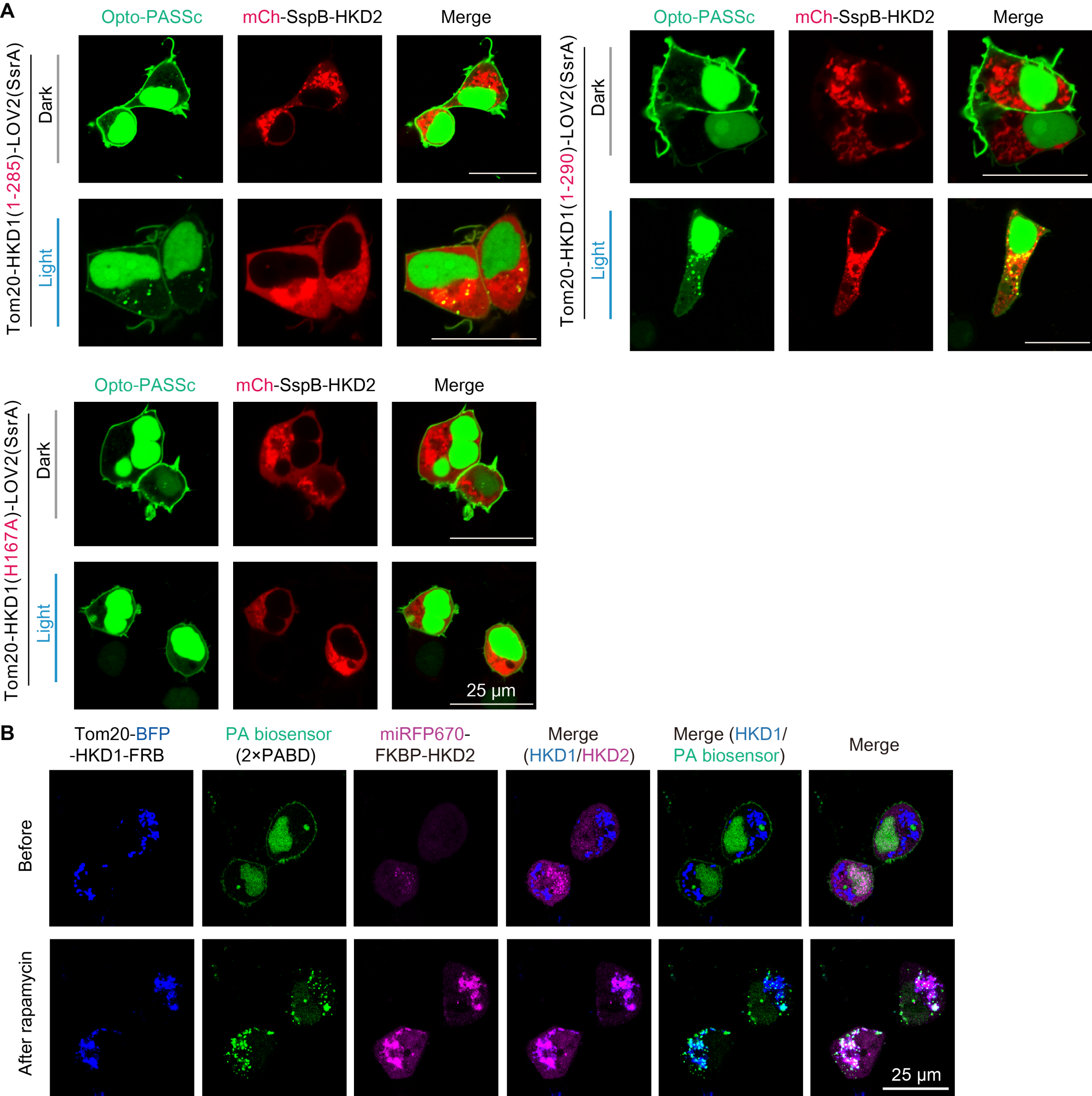

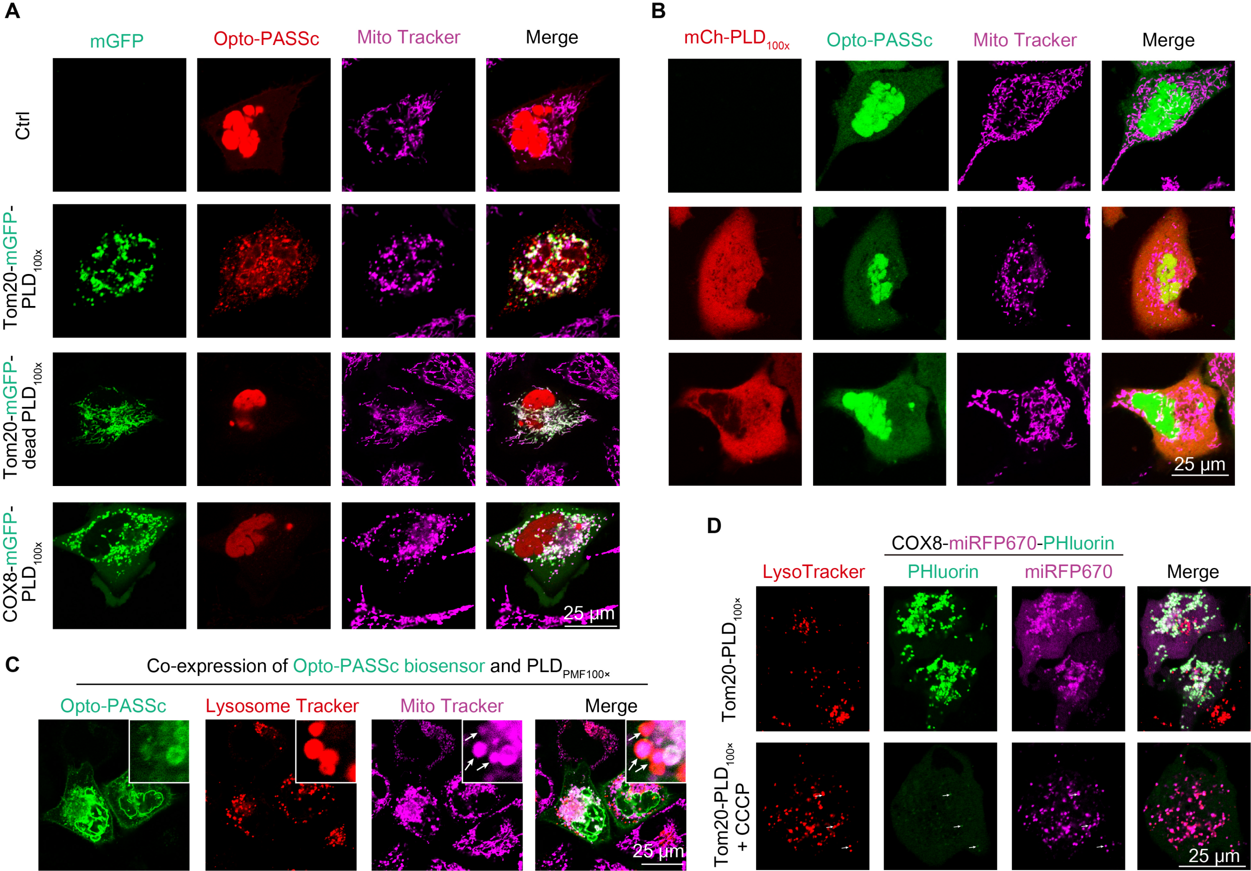
